# Chemical defense acquired via pharmacophagy can lead to herd protection in a sawfly

**DOI:** 10.1101/2022.01.14.476300

**Authors:** Pragya Singh, Neil Grone, Lisa Johanna Tewes, Caroline Müller

**Affiliations:** Chemical Ecology, Bielefeld University, Universitätsstr. 25, 33615 Bielefeld, Germany

**Keywords:** Sequestration, pharmacophagy, automimicry, plant-insect interaction, phytochemicals, *Hierodula patellifera* (Mantidae), Hymenoptera

## Abstract

Predation is an important selection pressure acting on organisms, with organisms evolving diverse anti-predator strategies to combat it. One such widespread strategy is chemical defense in which organisms either synthesize or extrinsically acquire defensive chemicals. Little is known about the intraspecific transfer of such chemicals and if such chemicals acquired from conspecifics can also serve as defense against predation. Here, we used adults of the turnip sawfly, *Athalia rosae*, which can acquire *neo*-clerodane diterpenoids (‘clerodanoids’) *via* pharmacophagy after exposure to the plant, *Ajuga reptans.* We show that clerodanoid access mediates protection against predation by mantids for the sawflies, both in a no-choice feeding assay and a microcosm setup. Moreover, even indirect access to clerodanoids, via nibbling on conspecifics that had access to the plant, resulted in protection against predation albeit to a much lower degree than direct access. Furthermore, sawflies that had no direct access to clerodanoids were less consumed by mantids when they were grouped with conspecifics that had direct access. Most, but not all, of such initially undefended sawflies could acquire clerodanoids from conspecifics that had direct access to the plant, although in low quantities. Together our results demonstrate that clerodanoids serve as chemical defense that can be intraspecifically transferred. Moreover, the presence of chemically defended individuals in a group can confer protection onto conspecifics that had no direct access to clerodanoids, suggesting a ‘herd-protection’ effect.

## 1. Introduction

Predation is an important biotic factor that many organisms in the wild encounter. To defend themselves, organisms exhibit a wide diversity of anti-predator strategies (Edmunds, 1974; Eisner *et al.*, 2005; Bergen & Beldade, 2019; Rowland *et al.*, 2020). Chemical defense is one such anti-predator strategy that is widespread amongst organisms, ranging from microorganisms (Matz *et al.*, 2008) to multicellular organisms (Santos *et al.*, 2016; Pančić & Kiørboe, 2018; Sugiura, 2020). Such defensive chemicals can either be synthesized *de-novo* or acquired extrinsically, for example, from the host plant diet (Opitz & Müller, 2009; Erb & Robert, 2016; de Castro *et al.*, 2021). For example, the oleander aphid, *Aphis nerii*, sequesters cardenolides from its host plant species, and utilizes these defensive compounds against both vertebrate and invertebrate predators (Züst *et al.*, 2018). Alternatively, organisms can specifically take up defensive chemicals independently of nutritional requirements, e.g. *via* pharmacophagy (Boppré, 1984; Nishida, 2014; Paul *et al.*, 2021b). For example, adults of some danaine butterfly species actively incorporate defensive chemicals like pyrrolizidine alkaloids from sources such as dried plant parts (Lawson *et al.*, 2021; Tea *et al.*, 2021). While these acquired chemicals confer protection on the individual taking them up, it is less well-elucidated whether and how this protection can extend to conspecifics that may not have access to these chemicals directly from the source.

The possibility that chemically defended individuals confer protection from predation on initially undefended conspecifics, what we coin as ‘herd-protection’, could be realized *via* different means. For example, individuals could acquire such chemicals not only directly from the (plant) source but also indirectly via intraspecific (Lawson *et al.*, 2021; Paul *et al.*, 2021a) or interspecific (Hashimoto & Hayashi, 2014; Tea *et al.*, 2021) interactions. Such indirectly acquired chemicals could then be used in the context of defense against predation.

Alternatively, after attacking a chemically defended individual, a predator may be deterred from attacking even chemically undefended conspecifics if it associates the phenotype with chemical defense by learned aversion or avoidance (Berenbaum & Miliczky, 1984; Hämäläinen *et al.*, 2020; Tuttle *et al.*, 2021). This phenomenon is often seen in combination with automimicry, wherein undefended individuals (mimics) benefit from the unpalatability of defended individuals (models) (Brower *et al.*, 1970; Aubier *et al.*, 2017). Furthermore, unpalatability or unprofitability of organisms is often associated with bright or aposematic coloration that functions as warning signal to predators (Cyriac & Kodandaramaiah, 2019; Kikuchi *et al.*, 2021). There is usually positive density-dependence in aposematism, such that conspicuous warning signals are more effective when they are common (Chouteau *et al.*, 2016; Kikuchi *et al.*, 2021).

Understanding how the presence of chemically defended individuals affects the rest of the population is an important question as studies have shown that there can be intraspecific variation in chemical defense, with some individuals lacking chemical defenses entirely (Best *et al.*, 2018; Prudic *et al.*, 2019; Sculfort *et al.*, 2020; Mattila *et al.*, 2021). Such variation could arise if the defensive chemicals have an associated cost, for example, for acquisition and/or maintenance of the chemical defense (Dimarco & Fordyce, 2017). Variation may also be influenced by intrinsic factors such as the age, sex, reproductive phase or immunological status of the individuals (Smilanich *et al.*, 2009; Zvereva & Kozlov, 2015; Arias *et al.*, 2016). In such scenarios with intraspecific variation in chemical defense, the degree of protection from predation for chemically undefended individuals depends on the frequency of defended and non-defended individuals (Gamberale-Stille & Guilford, 2004; Finkbeiner *et al.*, 2018). At higher density of chemically defended individuals, the motivation of predators could be reduced to search for undefended prey (Skelhorn *et al.*, 2011), it may be harder to detect undefended prey (Gamberale-Stille & Guilford, 2004; Skelhorn *et al.*, 2011), or more undefended individuals could indirectly access the chemicals and gain protection.

An excellent model system for examining the effect of defensive chemicals in deterring predation both directly and indirectly is the turnip sawfly, *Athalia rosae* (Hymenoptera: Tenthredinidae). The larvae of this species are well-studied for sequestering metabolites, i.e. glucosinolates, of their Brassicaceae host plants, which act as defense against various invertebrate and vertebrate predators (Müller & Brakefield, 2003; Müller & Arand, 2007; Opitz *et al.*, 2010; Matsubara & Sugiura, 2017). The bright orange adults still contain glucosinolates sequestered by the larvae (Müller & Sieling, 2006) but do not seem to be protected by these compounds against predators such as birds and lizards (Vlieger *et al.*, 2004; Boevé & Müller, 2005). However, *A. rosae* adults can additionally acquire other specialized metabolites, *neo*-clerodane diterpenoids (potentially together with other compounds, hereafter called clerodanoids), by pharmacophagy from certain plant species, such as *Ajuga reptans* or *Clerodendrum trichotomum* (both Lamiaceae) (Kawai *et al.*, 1998). Some of the clerodanoids found in insect bodies are likely slightly modified metabolic products from plant derived clerodanoids (Amano *et al.*, 1999; Paul *et al.*, 2021a). Moreover, these compounds can be acquired indirectly by nibbling on conspecifics that were exposed to plant material (Paul *et al.*, 2021a). While effects of clerodanoids on mating behavior have been shown previously (Amano *et al.*, 1999; Paul & Müller, 2021), empirical evidence for other functions such as in defense against predation is scarce and indirect (Nishida & Fukami, 1990). In the laboratory, the cultures are usually maintained without *A. reptans* leaves, suggesting that clerodanoid access is not essential for the sawflies’ survival (Paul *et al.*, 2021b, 2021a; Paul & Müller, 2021). Moreover, there is evidence for associated costs of clerodanoid uptake in *A. rosae*, such that adults with clerodanoid access exhibit a reduced lifespan (Zanchi *et al.*, 2021). Thus, most likely there is intraspecific variation in clerodanoid status by *A. rosae* in the wild.

We aimed to study whether uptake of clerodanoids can function as defense against predation both for focal individuals that directly acquire these compounds after access to plants, and indirectly for other conspecifics that come into contact with the focal individuals. Therefore, we observed the response of mantid predators in no-choice feeding assays to sawflies that either had access to clerodanoids directly from plants or indirectly from conspecifics or had no access (experiment 1). Next, we investigated survivorship of sawflies with or without clerodanoid access in both presence and absence of a predator (experiment 2). Lastly, we investigated if the presence of sawflies with clerodanoid access conferred protection from predation on conspecifics with no clerodanoid access and if this varied with their relative abundance (experiment 3). We also investigated the clerodanoid acquisition of sawflies from different treatments using chemical analysis (experiment 3). We predicted that both direct and indirect clerodanoid acquisition should lead to protection from predation by the mantids. Furthermore, presence of chemically defended sawflies should confer protection from predation on conspecifics, with protection increasing with proportion of chemically defended individuals in the group. We also expect sawflies without clerodanoid access that were grouped with conspecifics with clerodanoid access to acquire clerodanoids.

## 2. Materials and Methods

### (a) Experimental animals

The individuals of *A. rosae* used in this experiment were taken from a laboratory culture established using adults collected in the surroundings of Bielefeld, Germany, and supplemented annually with field-caught insects. The culture was maintained in mesh cages (60 × 60 × 60 cm) with overhead lighting in a laboratory at room temperature with a 16 h: 8 h light: dark cycle and ~60% relative humidity. Multiple females and males were put in a cage and provided with *Sinapis alba* (Brassicaceae) plants for the females to oviposit. Emerging larvae were raised on *Brassica rapa* var. *pekinensis* (Brassicaceae) plants. Males and females were collected and separated within two days of pupal eclosion. Adults were kept in a climate chamber at 20 °C (16 h: 8 h light: dark cycle, 70 % relative humidity) before being used in experiment 1 and 2, and were kept in the laboratory under culture conditions for experiment All adults were fed a maintenance diet of a honey-water mixture (1:50 dilution). Adults were allocated randomly to the different treatments of no clerodanoid access (C-), direct clerodanoid access (C+) or indirect clerodanoid access (AC+). For the C+ treatment, adults got access to a leaf section (0.8 cm^2^) of *A. reptans* for 48 hours, while for C-no *A. reptans* leaf was provided. For the AC+ treatment, adults got access to a C+ conspecific of the same sex (prepared as above) for 48 hours. *S. alba* plants were grown from seeds in a climate chamber (20 °C, 16 h: 8 h light: dark, 70% r.h.), while *B. rapa* and *A. reptans* plants were grown from seeds in a greenhouse (≥ 20 °C, 16 h: 8 h light: dark, 70% r.h.).

Twenty-three sixth instar individuals of *Hierodula patellifera* (Mantidae) were purchased (www.interaquaristik.de) and reared in individual cages (20 cm × 20 cm × 20 cm) on a diet of crickets in a climatized room (~20 °C) on a diet of crickets. The mantids used in the experiment were not exposed to the sawflies before experiment 1, and were starved 48 hours before experiment 1. Each mantid was offered a cricket before experiment 2 and 3 to avoid starvation effects over the experimental days, and those individuals that did not consume the cricket were excluded as mantids usually stop eating when nearing a molt.

### (b) Experiment 1: No-choice feeding assay to examine effect of clerodanoid access on predator deterrence

Mantids were placed individually in transparent containers (9.5 cm diameter, 20 cm height), one C-, C+ or AC+ female sawfly was introduced in the container (figure 1*a*), and the mantid response was noted. Each assay was conducted for 15 minutes unless the sawfly was consumed before this, at which point the assay was terminated. The predator response variables examined during the assay were number of attacks on sawfly, whether sawfly was discarded after mouth contact (SI 1a,b), and whether sawfly was consumed (SI 1c).

**Figure 1.**
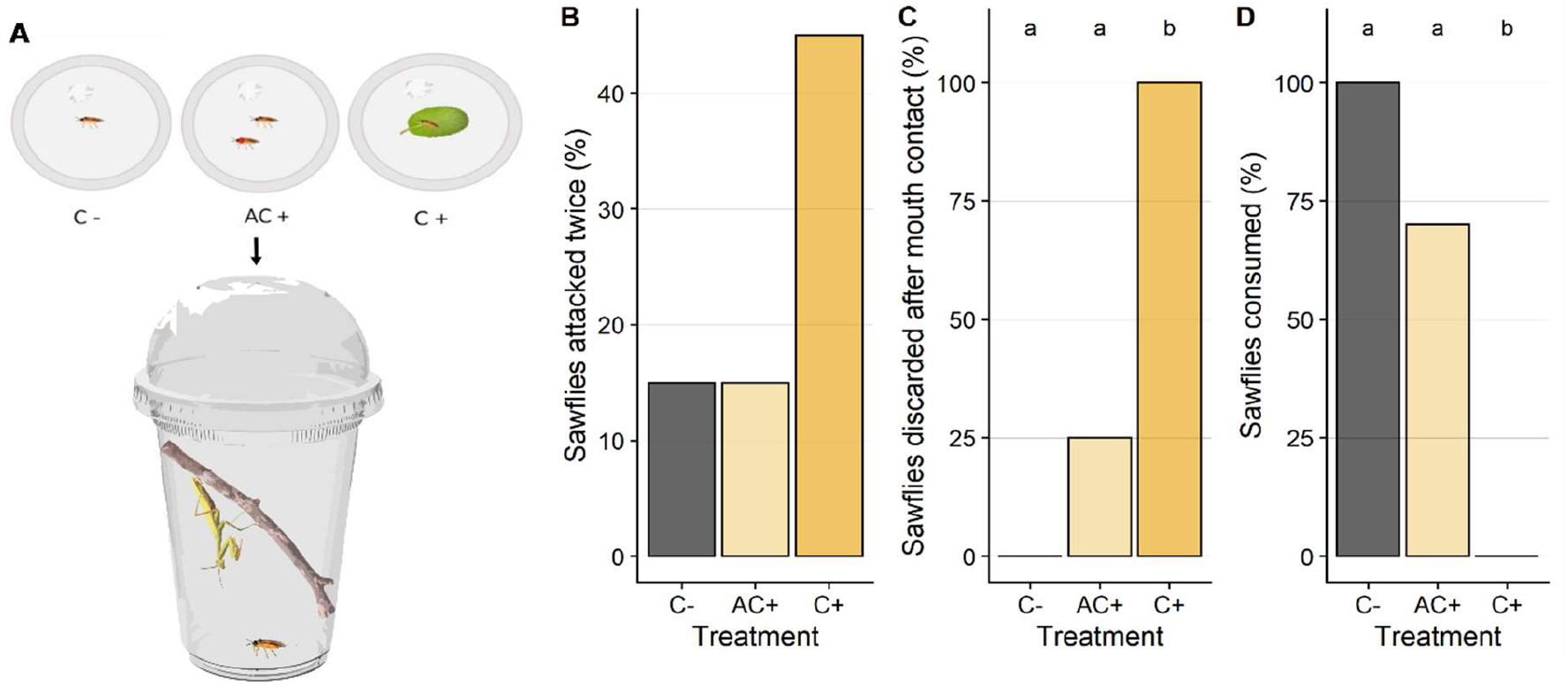
(A) Experimental design illustration of no-choice feeding assay, where each mantid was exposed to one *Athalia rosae* sawfly of different clerodanoid treatments (C-no access, AC+ indirect access via conspecific that had contact with leaf of *Ajuga reptans*, C+ direct access to *A. reptans*) over multiple trials performed in different orders. Effects of clerodanoid treatment on percentage of sawflies that were (B) attacked twice (all other sawflies were attacked once), (C) discarded after mouth contact, and (D) consumed, by mantid (*n* = 20 replicates per treatment). Different letters denote significantly different treatment effects inferred from Tukey HSD post hoc tests.

Discarding after mouth contact was a distinct behavior by the mantid, and could clearly be distinguished from the sawfly slipping or escaping from the mantid’s grab. For individuals that were attacked multiple times, we noted discarding as yes if the individual was discarded after mouth contact even once.

Sawflies of all treatments were offered to all mantids but in different orders; six mantids received *A. rosae* in the order C+, AC+, C-, seven in the order C-, C+, AC+, and seven in the order AC+, C-, C+ (total number of replicates N = 20). Upon noticing the sawfly, the mantids would orient for an attack. However, we did not compare the latency until attack, because the position of the mantids and how readily they noticed the sawflies differed between replicates. Two mantids did not attack sawflies of any of the treatments during the assay and were thus excluded from analysis as these mantids afterwards molted. We did not examine the long-term survivorship of sawflies that were discarded after attack by mantids, but damage to the sawfly spanned the spectrum from none to lethal (SI 1a-c).

We expected mantids to attack C-sawflies but not AC+ or C+ sawflies, e.g., if there were any repellent olfactory cues associated with uptake of clerodanoids by the sawflies. If AC+ or C+ individuals were attacked, we expected the mantids to discard the sawflies after tasting deterrent compounds and thus not consume them.

### (c) Experiment 2: Microcosm experiment to investigate clerodanoid access effect on survivorship in predator presence or absence

We used a fully factorial design (clerodanoid access × mantid presence) to evaluate the effect of clerodanoid acquisition on sawfly survival in the presence of a mantid predator in a microcosm (figure 2*a*). The four treatments were C-sawfly without mantid (C-M-), C-sawfly with mantid (C-M+), C+ sawfly without mantid (C+M-) and C+ sawfly with mantid (C+M+) with a sample size of eleven, ten, ten, and eight, respectively. All trials were performed in microcosm cages (25 cm diameter, 26 cm height) with a honey-water supply. The cages were kept in a climate room at 20 °C (16 h: 8 h light: dark cycle, 70% relative humidity) for the duration of the experiment. For each trial, we used five sawflies (three females and two males) and one or no mantid. We counted the number of sawflies alive every half-day for three days. For the mantid present trials, we also counted the number of sawflies ‘dead but not consumed’ at the end of three days, as sawflies may be attacked but not necessarily always completely consumed, e.g. if they are unpalatable. For these replicates, we calculated the number of consumed individuals as the difference between initial number of sawflies and the number of sawflies alive or ‘dead but not consumed’. This experiment was conducted two weeks after experiment 1 and the mantids were fed a cricket diet in the meantime. We expected C- with mantids to have reduced survival than sawflies in the other treatments. Moreover, we expected more C+ sawflies to be consumed by the mantids with time, which could be either due to a decreasing concentration of the clerodanoids or to prolonged starvation of the mantids.

**Figure 2.**
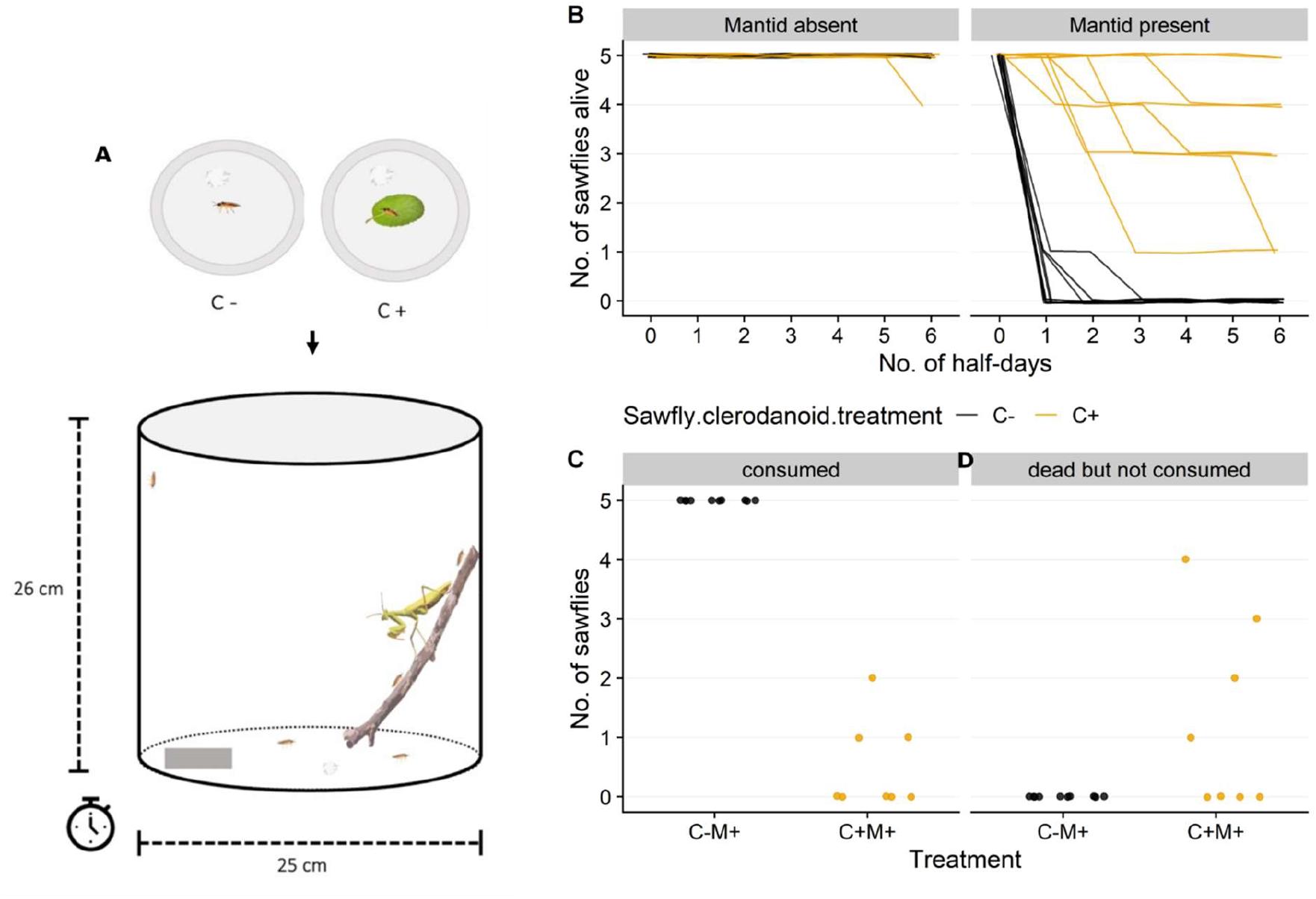
(A) Experimental design illustration of clerodanoid treatment (C-no access, C+ access to leaf of *Ajuga reptans*) and predation microcosm experiment, where in each microcosm five *Athalia rosae* sawflies were added that were either C- or C+ and with-or-without a mantid. B) Number of alive sawflies of different clerodanoid treatments over time. Lines are jittered to decrease overlapping. Number of C- and C+ individuals that were (C) consumed or (D) ‘dead but not consumed’ in replicates of mantid present (M+) treatment.

### (d) Experiment 3: Predation on C-conspecifics in mixed groups of C+ and C-sawflies in microcosm

We aimed to test whether presence of C+ sawflies led to defense against predation also for C-sawflies. Therefore, we set up four group-composition treatments, consisting of varying relative abundance of C+ and C-sawflies, keeping the total abundance of sawflies fixed at six. We chose six as the total abundance as we knew from experiment 2 that mantids can consume up to 5 C-*A. rosae* adults over a period of 1-3 half-days. The first group-composition treatment consisted of six C-sawflies (6C-), and the other three mixed group-compositions treatments were: two C+ and four C-sawflies (2C+4C-), three C+ and three C-sawflies (3C+3C-), and four C+ and two C-sawflies (4C+2C-). For each replicate, we joined the sawflies together in one petri dish according to the assigned group treatment two hours prior to adding them to the microcosm, to allow the sawflies to interact (e.g., mate or nibble).

We used eighteen mantids for the experiment, and a microcosm cage set-up identical to experiment 2. Each mantid was exposed to all group treatments in random order over multiple trials (four trials per mantid), with each trial lasting two days. Each mantid was fed a small cricket prior to each trial. For the mixed group-compositions treatments containing both C+ and C-individuals, we used only females as C- and males as C+. Using the different sexes allowed us to distinguish between C- and C+ individuals in each replicate. From experiment 2, we know that both male and female sawflies are consumed by mantids (see Results). In *A. rosae* adults, males are usually smaller than females (Sawa *et al.*, 1989; Travers-Martin & Müller, 2008), making it potentially easier for C-females to nibble and gain access to clerodanoids from C+ males. At the end of each trial, we counted the number of alive or ‘dead but not consumed’ C+ and C-sawflies for each replicate. We calculated the number of consumed C-sawflies as the difference between initial number of C-sawflies and number of C-sawflies alive or ‘dead but not consumed’. Similarly, we calculated the number of consumed C+ sawflies. We hypothesized that the presence and increasing abundance of C+ sawflies should lead to protection from predation for C-sawflies.

### (e) Chemical analysis of sawflies from experiment 3

To test whether C-sawflies in mixed group-composition treatments acquired clerodanoids compared to 6C-replicates, we collected C-sawflies from different group-composition treatments and analysed them chemically. We also collected C+ sawflies to confirm their clerodanoid acquisition. Lastly, we examined if C+ sawflies differed from C-sawflies of mixed group-composition treatment in the amount of clerodanoid acquired, i.e. if there is a difference between amount acquired directly from plant leaves or indirectly from conspecifics. We only collected individuals from replicates that had intact sawflies left at the end of the trial and stored them at −80 °C until further analysis. The final sample size for C-sawflies chemically analysed was six, five, seven, and five samples each of 6C-, 2C+4C-, 3C+3C- and 4C+2C-group-composition treatments, respectively. The final sample size for C+ sawflies was one, four and five from 2C+4C-, 3C+3C- and 4C+2C-group treatments, respectively. To extract putative clerodanoids from the sawflies, the individuals were freeze-dried and then homogenized using glass beads in a ball mill. Each individual was extracted twice by shaking for seven minutes at room temperature in ethyl acetate (LC-MS grade, VWR, Leuven, Belgium), and the supernatants were pooled to a final volume of 400 μl. The extracts were dried in a vacuum centrifuge at 35 °C. Dried extracts were suspended in 125 μl 100% methanol (LC-MS grade, Fisher Scientific, Loughborough, UK) in an ultrasonic bath for 15 minutes, after which they were filtered using syringe filters (polytetrafluoroethylene membrane, 0.2 μm pore size, Phenomenex, Torrance, CA, USA). The samples were analyzed using ultra high performance liquid chromatography (UHPLC; Dionex UltiMate 3000, Thermo Fisher Scientific, San José, CA, USA) with a Kinetex XB-C18 column (1.7 μm, 150 × 2.1 mm, with guard column, Phenomenex), and coupled to a quadrupole time of flight mass spectrometer (QTOF-MS; compact, Bruker Daltonics, Bremen, Germany), see SI 2 for details. The resulting chromatograms were processed with the software Compass Data Analysis 4.4 (Bruker Daltonics). The putative clerodanoids 482.22 m/z (C_24_H_34_O_10_) and 484.23 m/z (C_24_H_36_O_10_) occur in the chromatograms as [M+HCOOH-H]^−^ adducts resulting in features with 527.21 m/z and 529.23 m/z, respectively (Paul *et al.*, 2021a). We manually integrated the peak areas of these two features from the extracted ion chromatograms, extracted with 0.02 m/z accuracy.

We expected C-sawflies from mixed group-composition treatments to acquire clerodanoids, but to have lower amounts of clerodanoid compounds compared to C+ sawflies.

### (f) Statistical analyses

In experiment 1, we examined whether treatment had an effect on number of attacks, sawfly being discarded after mouth contact, and sawfly being consumed using a binomial generalized linear mixed-effects model (GLMM), with mantid identity as random effect. As there was quasi-complete separation in our data for two variables (sawfly being discarded after mouth contact and sawfly being consumed), i.e. the predictor variable could perfectly predict the response variable for a subset of our data, we fitted a Bayesian binomial GLMM using ‘blme’ package (version 1.0-5, Chung *et al.*, 2013) for all variables. We evaluated if the order in which the mantid was presented the treatments influenced the mantid response variables by incorporating the trial round of a treatment as a fixed effect (due to only three levels) in each model, and this gave qualitatively similar results (not shown).

In experiment 2, we examined the effect of treatment, time and their two-way interaction on number of alive sawflies using a Poisson GLMM (‘lme4’ package version 1.1-27.1, Bates *et al.*, 2015) with replicate ID as random effect. We also examined if there was a significant difference in number of consumed and ‘dead but not consumed’ sawflies between C- and C+ treatments with mantids (C-M+ and C+M+) using a Kruskal-Wallis test.

In experiment 3, we calculated the proportion of consumed, alive and ‘dead but not consumed’ C- with respect to the initial abundance of C-sawflies in that replicate. We examined whether the proportion of consumed, alive, and ‘dead but not consumed’ C-individuals differed significantly among the group-composition treatments using a binomial GLMM (‘lme4’ package), with mantid identity and trial number as random effects. We considered the number of C-individuals consumed/alive/‘dead but not consumed’ as successes and initial number of C-present as the number of trials. Similarly, we calculated the proportion of consumed, alive, and ‘dead but not consumed’ C+ with respect to the initial abundance of C+ sawflies in that replicate. Finally, we examined if there was a significant difference in the amount (quantified based on peak area) of the two putative clerodanoid compounds between C- and C+ sawflies of mixed group-composition treatments using a Kruskal-Wallis test.

All data was analyzed using R 4.0.5 (2021-03-31) (R Core Team, 2021). We checked and tested model assumptions statistically and visually. Posthoc-tests were conducted using ‘multcomp’ package (version 1.4-17, Hothorn *et al.*, 2008).

## 3. Results

### (a) Experiment 1. Clerodanoid access protects against consumption by predator

All sawflies irrespective of treatment were attacked at least once with no significant effect of treatment on number of attacks (*χ^2^* = 5.92, d.f. = 2, *p* = 0.052; figure 1*b*). In contrast, the treatments differed significantly in whether a sawfly was discarded after mouth contact by a mantid (*χ*^2^ = 21.15, d.f. = 2, *p* < 0.001; figure 1*c*, SI 3a), with C+ individuals dropped significantly more often compared to C-(posthoc test: *p* < 0.001) and AC+ (posthoc test: *p* < 0.001). Likewise, treatment had a significant effect on whether an individual was consumed (*χ*^2^ = 21.28, d.f. = 2, *p* < 0.001; figure 1*d*, SI 3b), with C+ individuals (0% consumed) being significantly less likely to be consumed compared to C-(100% consumed) (posthoc test: *p* < 0.001) and AC+ (70% consumed) (posthoc test: *p* = 0.001).

### (b) Experiment 2. C+ individuals are not consumed by predator even after prolonged exposure

There was a significant interactive effect of treatment and time on number of sawflies alive (*χ*^2^ = 36.76, d.f. = 3, *p* < 0.001). In the treatment with no mantids, all C- and all but one C+ individual survived across all replicates (figure 2*b*). In C-treatments with mantids, no individual was alive after three-half days. In contrast, for C+ treatments with mantids, all individuals were alive in two replicates. In the other six C+M+ replicates the number of alive individuals decreased with time, although in no replicate all individuals were killed. There was a significant difference in number of consumed sawflies (*χ*^2^ = 15.63, d.f. = 1, *p* < 0.001, figure 2*c*) with all sawflies consumed in the C-M+ treatment, while only few sawflies were consumed across three replicates in the C+M+ treatment. Similarly, the number of ‘dead but not consumed’ individuals significantly differed between C-M+ and C+M+ treatments (*χ*^2^ = 5.95, d.f. = 1, *p* = 0.014, figure 2*d*). In four C+ replicates but zero C-replicates, we collected ‘dead but not consumed’ individuals, suggesting that while the mantids attacked C+ individuals, they were not always consumed.

### (c) Experiment 3. Less C-individuals consumed by predator, if C+ individuals are present

Group-composition treatment had a significant effect on the proportion of consumed C-sawflies (*χ*^2^ = 41.87, d.f. = 3, *p* < 0.001; figure 3*a*), with significantly more C-consumed in the 6C-treatment compared to the 2C+4C-(posthoc test: *p* < 0.001), 3C+3C-(posthoc test: *p* < 0.001), and 4C+2C-(posthoc test: *p* < 0.001) treatments. There was no significant difference between the other treatments (SI 4a). The proportion of alive C-sawflies differed between the treatments (*χ*^2^ = 27.23, d.f. = 3, *p* < 0.001; figure 3*b*), with significantly less C-alive in the 6C-compared to the 2C+4C-(posthoc test: *p* = 0.005) and 3C+3C-(posthoc test: *p* < 0.001) treatment, while other treatments were not significantly different (SI 4b). Similarly, the proportion of ‘dead but not consumed’ C-sawflies differed between the treatments (*χ*^2^ = 8.77, d.f. = 3, *p* = 0.032; figure 3*c*), with significantly more C-sawflies being ‘dead but not consumed’ in the 4C+2C-compared to the 6C-treatment (posthoc test: *p* = 0.029), and no other significant differences between treatments (SI 4c). Similarly, to experiment 2, C+ sawflies were consumed rarely, although in many replicates there were ‘dead but not consumed’ C+ individuals (SI 5a-c).

**Figure 3.**
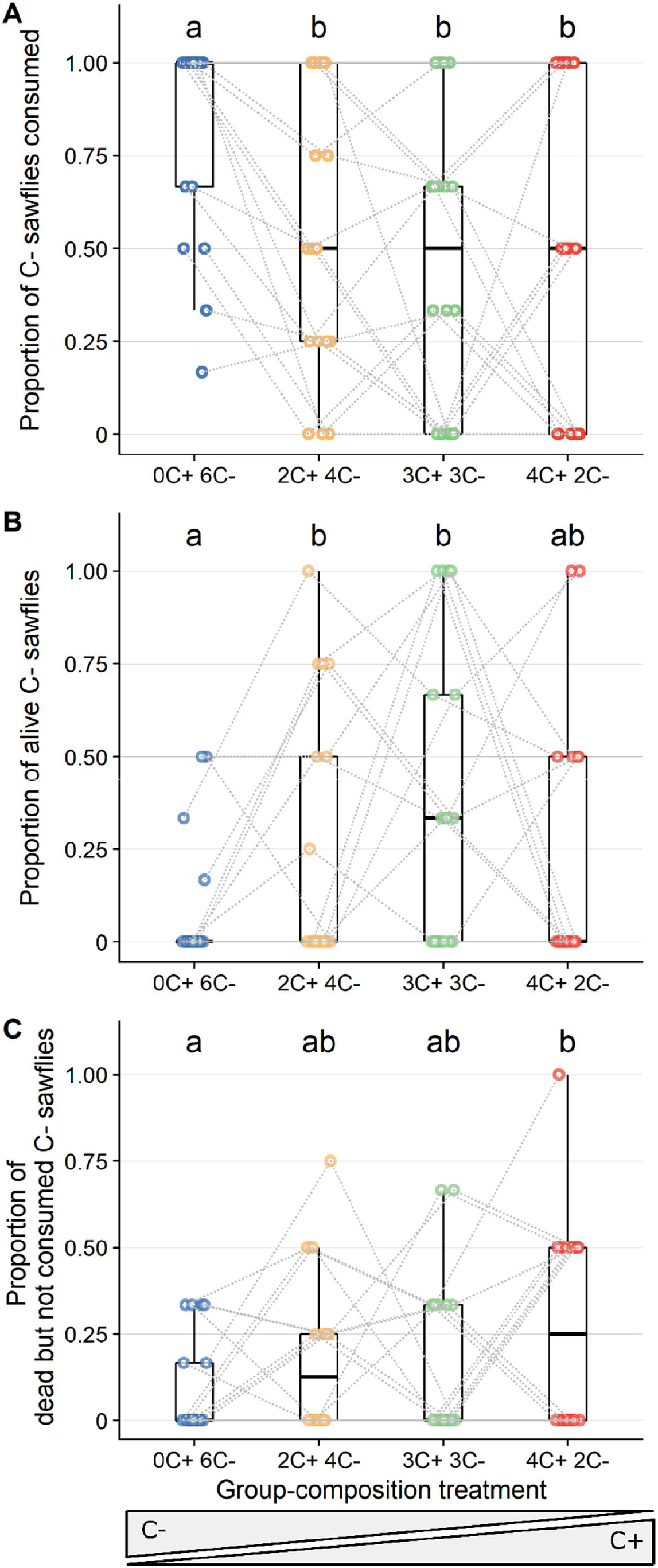
Effects of different group-composition, i.e. varying relative abundance of *Athalia rosae* sawflies with (C+) and without (C-) access to a leaf of *Ajuga reptans* and thus clerodanoids, on proportions of (A) consumed, (B) alive, and (C) ‘dead but not consumed’ C-sawflies in presence of a mantid. Data are presented as boxplots, medians and interquartile with individual data points also plotted. Grey dotted lines connect data from each mantid (*n* = 18) across trials. Note that abundance of C-sawflies decreases and C+ increases from left to right.

The chemical analysis revealed that the putative clerodanoids 482.22 m/z and 484.23 m/z could be detected in sixteen (~94%) and fourteen (~82%), respectively, of the seventeen sampled C-sawflies from mixed group-composition treatments (figure 4*a,b*). All ten (100%) C+ sawflies had acquired both putative clerodanoids (figure 4). There was intraspecific variation in the amount of clerodanoids acquired for both C+ and C-sawflies of mixed groups. Two out of six replicates of the 6C-group-treatment also had small amounts of clerodanoids (figure 4), possibly resulting from contamination as these replicates were placed in the microcosms where previously mixed group-composition treatments were housed. This contamination could also potentially explain why these C-sawflies were not consumed by the mantids. C+ sawflies had significantly higher amounts of both clerodanoids, 482.22 m/z (*χ*^2^ = 10.98, d.f. = 1, *p* < 0.001, figure 4*a*) and 484.23 m/z (*χ*^2^ = 15.75, d.f. = 1, *p* < 0.001, figure 4*b*), than C-sawflies for mixed group-composition treatments.

**Figure 4.**
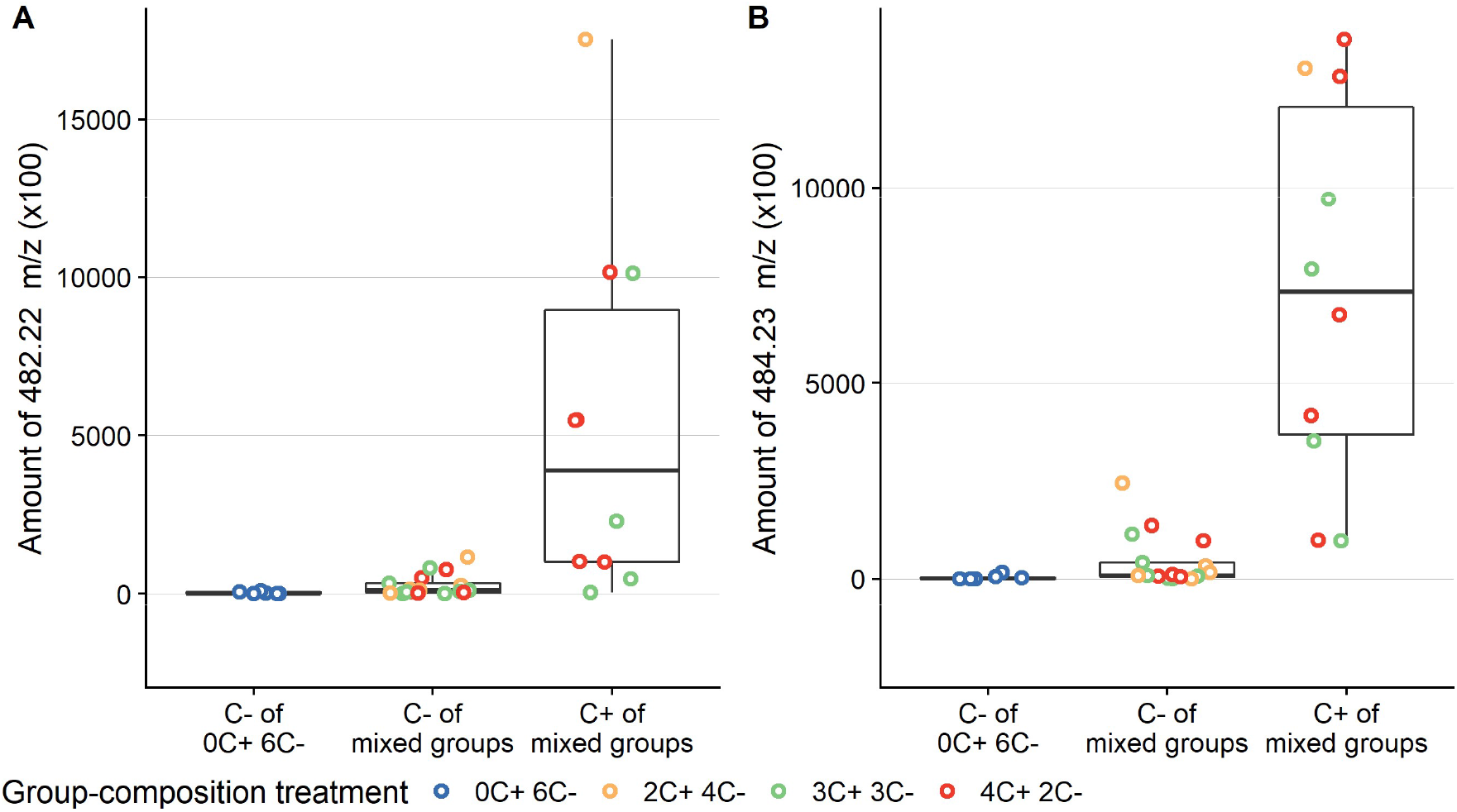
Amount (peak area) of candidate chemical features representing putative clerodanoids, (A) 482.22 m/z (C_24_H_34_O_10_) and (B) 484.23 m/z (C_24_H_36_O_10_), respectively, from the extracted ion chromatograms for C-(without access to a leaf of *Ajuga reptans*) and C+ (with access to a leaf of *A. reptans*) sawflies of different group-composition treatments. Mixed groups represent groups that had both C+ and C-sawflies present in the microcosm, while 0C+6C-had only C-sawflies present.

## 4. Discussion

Chemical defense as an anti-predator strategy is widespread and well-documented in animals (Speed *et al.*, 2012). However, it is less clear if such chemical defense can be intraspecifically transmitted and whether presence of chemically defended individuals can confer protection onto undefended conspecifics. In some species, such defensive chemicals can be taken up independently of the nutrient-delivering food source, as in the case of our study organism, *A. rosae*, that takes up clerodanoids (and potentially also other chemicals) pharmacophagously from *A. reptans* plants and likely metabolized them (Paul *et al.*, 2021b). Here, we showed that direct access to *A. reptans* leaves serves as defense against predation by making the sawflies unpalatable to the predator. We also demonstrated that clerodanoid access provides protection not only to focal individuals but also to conspecifics in mixed groups of C+ and C-individuals.

While most sawflies without access to clerodanoids were lethally attacked and consumed by the mantids, only very few sawflies that had taken up clerodanoids from leaves were consumed, as we had predicted. Nevertheless, many sawflies with clerodanoid access were also attacked but then readily rejected by the mantids, as shown by the ‘discarding after mouth contact’ behavior in experiment 1 and by the higher number of ‘dead but not consumed’ sawflies in experiment 2. This rejection is likely induced by the bitter taste of the clerodanoids that may be deposited on the cuticle and in body tissue of the adult sawfly (Nishida & Fukami, 1990). When tested directly, two clerodanoids (clerodendrin B and D) had a deterrent effect on Japanese tree sparrows, who consumed fewer rice grains that had been treated with the clerodanoids compared to untreated grains (Nishida & Fukami, 1990). A taste-rejection behavior, in which predators taste but do not ingest a prey item, as found by the mantids in the present experiment, has been shown to be elicited by distasteful prey and can lead to an increased survivorship of the prey (Halpin & Rowe, 2017). Although we did not quantify the long term survivorship of sawflies after mantid attack in experiment 1, visual inspection showed that the damage spectrum ranged from nearly unharmed to dead sawflies (SI 1a,b), indicating that clerodanoid uptake could lead to survivorship advantages.

Adult *A. rosae* can acquire clerodanoids not only from plants but also from conspecifics *via* nibbling on their body surface. However, acquiring clerodanoids indirectly from conspecifics resulted in less protection than direct acquisition from the plants in *A. rosae* in our experiments. Not all sawflies successfully acquire sufficient clerodanoid amounts from C+ conspecifics (Paul *et al.*, 2021a) and the concentrations could be much lower than after direct uptake from the *A. reptans* leaves (figure 4). Such quantitative and potentially also qualitative differences in clerodanoid acquisition may explain, why many AC+ sawflies (experiment 1) and C-sawflies in mixed C+ C-group treatments (experiment 3) were consumed by mantids. The effectiveness of defense chemicals sequestered from host plants against predators has been shown to be concentration-dependent in a glucosinolate-sequestering leaf beetle species, with individuals having lower levels of sequestered glucosinolates being more susceptible to predation (Sporer *et al.*, 2020). In *A. rosae*, transfer of clerodanoids can occur both within and between sexes (Paul *et al.*, 2021a), which seems rather exceptional. In other insect species, usually chemicals are transferred from the male to the female during mating. For example, in some arctiid moth species, pyrrolizidine alkaloids are sexually transmitted from males to females, and these chemicals render protection against predation to the recipient female (Gonzalez *et al.*, 1999; Conner *et al.*, 2000). Moreover, such defensive chemicals acquired by the female from the male can also be incorporated into the offspring (Eisner *et al.*, 2002; Camarano *et al.*, 2009; Sternberg *et al.*, 2015) providing it with benefits. Evidence for such benefits of parental clerodanoid access to offspring in *A. rosae* is currently lacking (Paul *et al.*, 2021b).

Our data from the predation microcosm experiment (experiment 2) demonstrated that the mantids attacked C+ sawflies, with number of alive sawflies decreasing over time, but this decline was less rapid than that of C-sawflies in presence of mantids. This suggests that the mantids may learn to avoid C+ sawflies after first encounters, leading to a longer survival period of the sawflies. Learned aversion has been found in the mantid *Tenodera aridifolia*, where the mantids reduced attacks on certain prey items when these prey items were made bitter (Carle *et al.*, 2018). Similarly, repeated exposure to unpalatable milkweed bugs that had sequestered cardenolides from their host plants led to avoidance by *T. aridifolia* of both palatable and unpalatable milkweed bugs altogether (Berenbaum & Miliczky, 1984).

Interestingly, the mantid *H. patellifera* attacked sawflies in both experiment 2 and 3 despite having been exposed to the C+ sawflies previously, suggesting that such avoidance learning may not last long, as also seen in *T. aridifolia* (Prudic *et al.*, 2007). Moreover, such avoidance learning can be more effective if the distasteful prey is conspicuously colored (Roper & Redston, 1987; Raška *et al.*, 2017). Indeed, organisms that use chemical defenses are usually, but not always, brightly colored and conspicuous, i.e. aposematic, to advertise their distastefulness to predators (Poulton, 1890; Kikuchi *et al.*, 2021). The sawfly *A. rosae*, used in our study, is aposematically colored, with a bright orange body (SI 1), which might facilitate temporary avoidance learning by mantids.

In line with our prediction, the presence of C+ sawflies was beneficial for C-sawflies with a smaller proportion of C-sawflies consumed in group-composition treatments that had C+ individuals present compared to groups of C-sawflies only. This suggests that presence of chemically defended sawflies can lead to ‘herd protection’ of conspecifics. Such protection may result from the C-individuals acquiring clerodanoids, or temporary learned avoidance of C- by mantids after encountering C+ sawflies. Our chemical analysis showed that most, but not all, sawflies had acquired detectable amounts of clerodanoids, suggesting that both of these mechanisms could play a role in the ‘herd-protection’ of C-conspecifics. In our experiment, there was no significant change in the expected direction in number of consumed C-sawflies across the gradient of the C+ and C-mixed group-composition treatments. This may have been due to the small number of sawflies used (six), and hence only small differences between the mixed treatments. A study using the domestic chick, *Gallus gallus domesticus*, as predator showed that the birds rejected the mimics (palatable prey items) less frequently when the relative abundance of these mimics compared to the models (unpalatable prey items) increased, but birds only discriminated between the models and mimics when the frequency of mimics was above 25% (Skelhorn & Rowe, 2007). Moreover, density-dependence of predation can also be influenced by other factors such as predator behavior, e.g. learning, forgetting and memory (Speed & Turner, 1999), and the energetic and informational state of the predator, leading to state-dependent decision-making (Aubier & Sherratt, 2020).

In natural populations of *A. rosae*, intraspecific variation in clerodanoid uptake could be expected if the distribution of pharmacophagy-suitable plants is patchy, if there in intraspecific variation in the clerodanoid concentrations available from the plants, or if there are associated costs of clerodanoid uptake. Indeed, *A. rosae* individuals that were exposed to clerodanoids had a shorter lifespan than control individuals (Zanchi *et al.*, 2021). This suggests that there might be costs to clerodanoid uptake, although clerodanoid access did not immediately cause high mortality during our observed period in experiment 2 (figure 2*b*). Similar costs of chemical defense have been revealed in swallowtail butterflies of the tribe Troidini which showed reduced larval survivorship (Dimarco & Fordyce, 2017) or a reduction in adult fat content (Fordyce & Nice, 2008) when sequestering toxic alkaloids. Our study demonstrates that even if individuals do not take up clerodanoids, they could still benefit indirectly from conspecifics that do. This could lead to emergence of cheaters that do not pay the cost of chemical defense but enjoy the benefits (Lindstedt *et al.*, 2018). Future studies should examine variation in clerodanoid contents in natural populations of *A. rosae*. In conclusion, our study showed that clerodanoids serve as chemical defense for *A. rosae* that can be intraspecifically transferred. Furthermore, chemically defended sawflies can confer protection onto conspecifics that had no direct access to clerodanoids in a group, indicating a ‘herd-protection’ effect.

## Acknowledgements

This study was funded by the German Research Foundation (DFG) as part of the SFB TRR 212 (NC^3^), project number 396777467 (granted to CM).

